# Abnormal behavioral episodes associated with sleep and quiescence in Octopus *insularis*: Possible nightmares in a cephalopod?

**DOI:** 10.1101/2023.05.11.540348

**Authors:** Eric A. Ramos, Mariam Steinblatt, Rachel Demsey, Diana Reiss, Marcelo O. Magnasco

**Author notes:** **Correspondence:** Eric Angel Ramos.

## Abstract

This paper presents some unusual behaviors observed in one single specimen of *O. insularis*. While nothing can be concluded rigorously from such data, we share the data and our analysis with the community, in the hope that others will be on the lookout for such rare events. Sleep is a fundamental biological function that is present in all tested vertebrates and most invertebrates.

Cephalopods, such as octopuses, are cognitively complex animals that display active and inactive sleep states similar to those of vertebrates. In particular, octopuses have active sleep states during which they display sequences of camouflage patterns and modulation of basal rhythms, while remaining relatively unresponsive to outside stimuli. Some scientists have speculated that these states could be analogous to dreaming in mammals, involving episodic recall with a narrative structure. The convergent evolution of sleep in neurologically complex animals is a striking possibility, but its demonstration requires overcoming significant challenges. Towards this end, capturing abnormal sleep-associated episodes and other parasomnias in cephalopods can provide further insight into the biology of their sleep. This study reports abnormal behavioral episodes associated with transitions between activity states and sleep states observed in a male *Octopus insularis*. The study used continuous video monitoring to characterize the animal’s activity patterns and detect rare behavioral episodes. Over the course of a month, four brief episodes (duration range: 44-290 seconds) were identified during which the octopus abruptly emerged from quiescent or active sleep, detached itself from its sleep position, and engaged in antipredator and predatory behaviors (with no predator present). The longest of these episodes resembled the species-typical response to a predatory attack, suggesting that the animal may have been responding to a negative episodic memory or exhibiting a form of parasomnia. These findings, in conjunction with recent evidence for sleep in octopuses, highlight the complexity of possible sleep-associated behavioral episodes. Investigating sleep in invertebrates is crucial to understanding the evolution of sleep across distantly related species.

## INTRODUCTION

Sleep is a fundamental physiological process and behavior present in most amniotes, such as mammals (Siegel, 2005), birds (Lesku & Rattenborg, 2014), and reptiles (Shein-Idelson et al., 2016). It is characterized by alternating states of neural activity and inactivity in electrophysiological recordings of amniotes (Noda et al., 1969), manifesting in cyclic shifts between periods of rapid eye movement sleep and non-REM sleep (Blumberg et al., 2020; Peever & Fuller, 2017). In vertebrates, sleep serves critical biological functions, including the reduction of oxidative stress, and synaptic consolidation of memory (Xie et al., 2013; Blanco et al., 2015). Sleep-like states have been documented in invertebrate species of different evolutionary lineages (Jha & Jha, 2020 for review) including *Drosophila melanogaster* (Shaw et al., 2000; Greenspan et al., 2001), crayfish (Ramon et al., 2004), *Caenorhabditis elegans* (Raizen et al., 2008), and jellyfish (*Cassiopea*; Nath et al., 2017). This indicates that analogues of sleep are evolutionarily conserved (Campbell & Toblew, 1984; Siegel, 2005; Corner, 2013; Keene & Duboue, 2018).

Cephalopods are intelligent, large-brained animals that display a range of complex cognitive functions, including long-term memory, learning, and the ability to discriminate complex stimuli (Hanlon & Messenger, 2017). The complex body patterning of cephalopods serves multiple functions, allowing dynamic camouflage in different environments necessary for predator evasion and possibly for communication with conspecifics (Adamo et al., 2006; Staudinger et al., 2013; Hanlon & Messenger, 2017). Additionally, cephalopods display a remarkably diverse and adaptive behavioral repertoire, evident in the complexity of their habitat use and predatory behaviors across their distribution range (Ambrose, 1984; Leite et al., 2009; Scheel et al., 2017).

Cephalopods including *Octopus insularis, O. vulgaris*, and the common cuttlefish (*Sepia officinalis*) are known to display rest-activity cycles with ultradian periodicities, similar to the sleep states of amniotes (Meisel et al., 2003; Meisel et al., 2011; Frank et al., 2012; Iglesias et al., 2017; Medeiros et al., 2021). In laboratory experiments with *O. vulgaris*, sleep deprivation caused compensatory sleep states supporting the presence of homeostatic sleep mechanisms (Brown et al., 2006). Similarly, *S. officinalis* exhibit sleep-like states in which they remain inert while displaying periodic color changes and arm twitching (Duntley et al., 2002; Duntley & Morrissey, 2004; Frank et al., 2012). Recent studies further identified the cyclic nature of REM-like sleep states alternating with a quiescent sleep-like state in an ultradian pattern in *O. insularis* (Medeiros et al., 2021). Observations included an active sleep state and sleep state with one eye movement, possibly analogous to rapid-eye-movement (REM) sleep (Medeiros et al., 2021). These sleep states have been also associated with brain activity in octopuses, supporting differences in neural activity in wake and sleep states akin to vertebrate sleep (Gutnick et al., 2023).

Dreaming is a further mystery within the evolutionary enigma of sleep. Dreams, as experienced by humans, are vivid hallucinatory episodes characterized by having a narrative structure (Aron et al., 1989; Hartmann, 1998). Understanding whether animals dream poses unique challenges, stemming from our inability to access their internal subjective experience. Cephalopod camouflage offers a unique way to access outputs of internal states inaccessible from other non-camouflaging species. Recently Malinowski et al. (2021) conjectured that the characteristic sequences of camouflage changes occurring during episodes of active sleep in a specimen of *O. cyanea* appear to have a narrative structure that follows a sequence of natural behaviors. The authors argue that these behaviors may represent dream-enactment in cephalopods, in which the dreamer actively acts out dream actions, driven by the possible linkages between their waking and sleep behaviors (Malinowski et al., 2021). They also argue that in the active sleep states, motor responses and body pattern displays are only partially suppressed. Alternatively, parasomnias are involuntary episodes associated with unintended motor and behavioral activity in the transition to and from sleep (Pavlova et al., 2015); dream disturbances and nightmares such as those associated with PTSD (Campbell & Germain, 2016) are typically correlated with sleep state transitions (Nielsen & Zadra, 2010). It is unknown if cephalopods display these behaviors. However, given cephalopods so closely match the behavioral definitions for sleep in vertebrates, classifying behavioral phenomena and their contexts may help to identify evolutionary parallels in sleep functions, stages of sleep, and transitions from sleep to waking states across taxa.

This study reports observations of abnormal sleep-associated behavioral episodes in one male *O. insularis*, in which the octopus displayed parasomnia episodes consistent with dream-enactment of nightmares in vertebrates (though not inconsistent with other potential explanations given the single-subject nature of this study). We contextualize this occurrence, provide descriptions of the behavioral sequences exhibited and discuss the implications for the convergent evolution of sleep mechanisms in vertebrates and cephalopods.

## MATERIALS AND METHODS

### Animal housing and care

This study was conducted at the Laboratory of Integrative Neuroscience and the Comparative Biosciences Center at The Rockefeller University. A live-captured male *Octopus insularis* was obtained from the coastal waters of the Florida Keys by a vendor in Marathon, Florida (Dynasty Marine Associates Inc.). Species was confirmed as O. insularis through genomic analysis and visual identification (Maloney et al., *In review*). The animal weighed 514.4 g at death and was identified as male through the presence of a hectocotylus on its third arm on its right. The animal arrived at the lab on 11 February 2020 and died naturally on 04 April 2020; given that many definitions of senescence state that it can be a months-long process, the subject would have satisfied that definition at the time of observations; however, there were no observable signs of senescence until a week before its death. All research protocols were approved by the Institutional Animal Care and Use Committee (IACUC) at The Rockefeller University. The welfare of the animal was the primary concern, thus the EU directive on cephalopod care (Directive 2010/63/EU, European Parliament and Council of the European Union, 2010) was followed and no direct harm was caused to the animal. The octopus was allowed to senesce naturally. Upon its death, a necropsy confirmed the presence of the parasitic coccidian protozoan of the genus *Aggregata* in the digestive tracts, a common finding in cephalopods that negatively impacts their well-being by weakening their immune function and destroying their intestinal mucosa and other digestive tissues (Pascual et al., 2010; Castellanos-Martínez et al., 2019).

It was housed individually in a 120-gallon glass tank (Aqueon, 121.9 cm length × 45.7 cm width × 71.1 cm height). Transparent plastic panes with a thin blue film were used to cover the rear and one side wall of the tank to create a sufficiently sheltered atmosphere for the animal. Sand covered the bottom of the tank to prevent viewing of the reflective surface of the bottom of the tank. To provide ample stimulation and enrichment for the animal, a variety of items were added throughout the tank over time, including Real Reef Rocks (Reel Reef Solutions); a small gray PVC tube (10 cm diameter × 30 cm length); a large gray PVC elbow joint (50 cm diameter); numerous small white PVC pieces and green artificial plants; and real seashells. Damselfish were co-housed to provide additional active stimulation.

The tank was connected to a closed circulation system for filtration centered in a 139.1 L three-partition sump (91.4 cm length × 34.6 cm width × 38.1 cm height) (Trigger 36 Crystal Sump) housed under the tank, connected via several tubes for filtration. Water was pumped through two inputs at the top of the animal tank through two rubber tubes into a 25-watt UV sterilizer (Emperor Smart) through filter socks in one sump partition. The second sump housed an in-sump protein skimmer (Reef Octopus Classic 150SSS) for extraction of organic waste. Filtered water was pumped back into the animal tank from the third sump partition with a water pump (Jecod/Jebao DCT-4000) via a single outflow at the top of the animal tank. Water was pumped at maximum to provide sufficient oxygenation for the octopus. Fluidized beds, activated charcoal, and a variety of biological filter media were added to the sump as a substrate for growth of bacterial colonies needed for the nitrogen cycle: consuming ammonia and converting it to nitrites and nitrates. A sandstone plate was connected to air compressors and placed at the bottom of one sump partition under the filter media to provide aeration and fluidization necessary for bacterial growth. Active bacteria were regularly added to increase bacterial populations. Water temperature was regulated with a heating element (Finnex Titanium) and a digital temperature controller (Aqua Logic).

Water quality was rigorously monitored and maintained through daily extraction of waste (e.g., sucker cuticles shed by the octopus, animal excrement, food remains) from filter socks and sand washes with a siphoning gravel washer concurrent with water changes of 15-20% of tank water. Water was replaced with reservoirs of a saltwater admixture made with deionized water and artificial reef salt (Instant Ocean, Blacksburg, VA). A variety of amphipods and copepods were added to improve tank biodiversity and health, and several species of detritivores were added to help breakdown and remove organic waste. An Apex Fusion controller system (Neptune Systems, Morgan Hill, CA) enabled continuous remote monitoring of pH, salinity, and water temperature with calibrated probes placed at the top of the water column in the animal tank. The system also regulated the automated 12 hr/12 hr light/dark cycle of the overhead light fixtures to simulate natural light conditions. Lighting consisted of LED light bulbs in conical aluminum light fixtures, two 250-watt bulbs provided white light in the daytime (0730–1930 h) and two 36-watt bulbs provided deep red light (660 nm) at night (1930–0730 h). Water quality parameters was manually tested multiple times daily for pH, salinity, water temperature, and the concentration of ammonia, nitrates, nitrates using API testing kits and handheld electronic sensors (Hanna Instruments, Woonsocket, RI) to maintain optimal levels for the animal (pH: 8.2–8.5; salinity: 32–34 ppt; water temperature: 23–25 C°; ammonia: 0 ppm; nitrates: <20 ppm; nitrites: 0 ppm).

The animal was fed 50–150 g pieces of thawed frozen shrimp or whole shrimp once to twice a day (1100 h and 1600 h). It was periodically fed live crabs for enrichment. Pieces of shrimp were either dropped near the animal or placed into various rubber dog toys (Kong), plastic jars, or clear acrylic tubes.

### Continuous video observations

Octopus activity was monitored 24 hours per day with a Lorex 4K Ultra HD 8 Channel Security System filming with 6 bullet cameras (8 MP, 105° field of view, 70 mm width × 180 mm length) routed to an 8-channel video recorder. Three cameras were positioned to capture different perspectives of the tank, filming continuously 24 hr/day (H.265 compression, 3840 × 2160 pixels; 15 fps). Cameras were connected by ethernet to the video recorder, enabling live viewing through a computer monitor and live streamed through the local Wi-Fi network. Footage was stored locally in the recorder in 30 min segments and was exported in .mp4 format or in .dav format (and later converted to .mp4) to a RAID physical drive.

### Video analysis

Five minutes after Episode 3, one of the authors (EAR) noticed and recorded the octopus’s abnormal behavior as it happened and then reviewed the security camera footage to document the entire episode. Three distinct characteristics were identified and used to identify similar episodes in the future: 1) sudden changes in the octopus’s body pattern and movement *while it was stationary or in a quiescent state*, 2) disordered movements of its arms and body, including seemingly involuntary contortions, and 3) a series of evasive and/or antipredator responses such as inking, using its posterior jet, and expelling water from its siphon.

To identify similar episodes, we reviewed our database of continuous security footage and high-resolution footage for instances involving these characteristics. Octopus behaviors were initially classified as ‘Active’, ‘Alert’, or ‘Quiet’ as defined by Brown et al. (2006). Quiescent behaviors were classified according to Medeiros et al. (2021) defined in Table 1 and illustrated in Figure 1 (see Supplementary Video S1 for examples). Quiet or quiescent states (Medeiros et al. 2021) were reviewed multiple times because normal sleep in the animal involved abrupt changes in body pattern and movement while stationary and/or quiescent but not in association with any other behaviors. We also identified and categorized the behavioral states preceding and following each episode to identify if these episodes occurred within sleep states or at transitions into and out of sleep states.

**Table 1.** Description of active and quiescence states used to characterize the normal and abnormal sleep behaviors in a male *Octopus insularis*. Definitions adapted from Brown et al. (2006) and Medeiros et al. (2021).

| Behavior state | Description | Source (s) |
| --- | --- | --- |
| Alert | Little to no movement of the body. Slight movements of the arms and body with eyes open. | Brown et al., (2006) |
| Active | Active movement of the entire body around its enclosure. | Brown et al., (2006) |
| Active sleep (quiet with dynamic body pattern and eye movements) | Dynamic skin color and texture changes. Both eyes in movement. Muscular twitches with contractions of suckers, arms, and body. Also named quiet with dynamic body pattern and eye movements. | Medeiros et al., (2021) |
| Quiet sleep/Quiescence (quiet with closed pupil) | Quiescent and largely motionless body with pale body color. Pupils closed and narrowed to a thin slit. Sporadic movements of suckers and arms with less intensity than in active sleep. Camouflaged when on a rock. | Medeiros et al., (2021) |
| Quiet/Quiescence with open pupil | Quiescent and largely motionless body with pale body color when in tube or on wall or camouflaged when on a rock. Sporadic contraction and expansion of arms. Regular respiration. | Medeiros et al., (2021) |
| Quiet with half and half body pattern | Animal motionless with abrupt color change to dark red. Colors are bilaterally symmetric with one side remaining pale and white or pale and slightly mottled. Sometimes involves exophthalmos (i.e., bulging and protruding of the eye) | Medeiros et al., (2021) |
| Quiet with only one eye movement | Cyclic movement of one eye including dilation, contraction, and exophthalmos (i.e., bulging and protruding of the eye). | Medeiros et al., (2021) |

**Figure 1.**
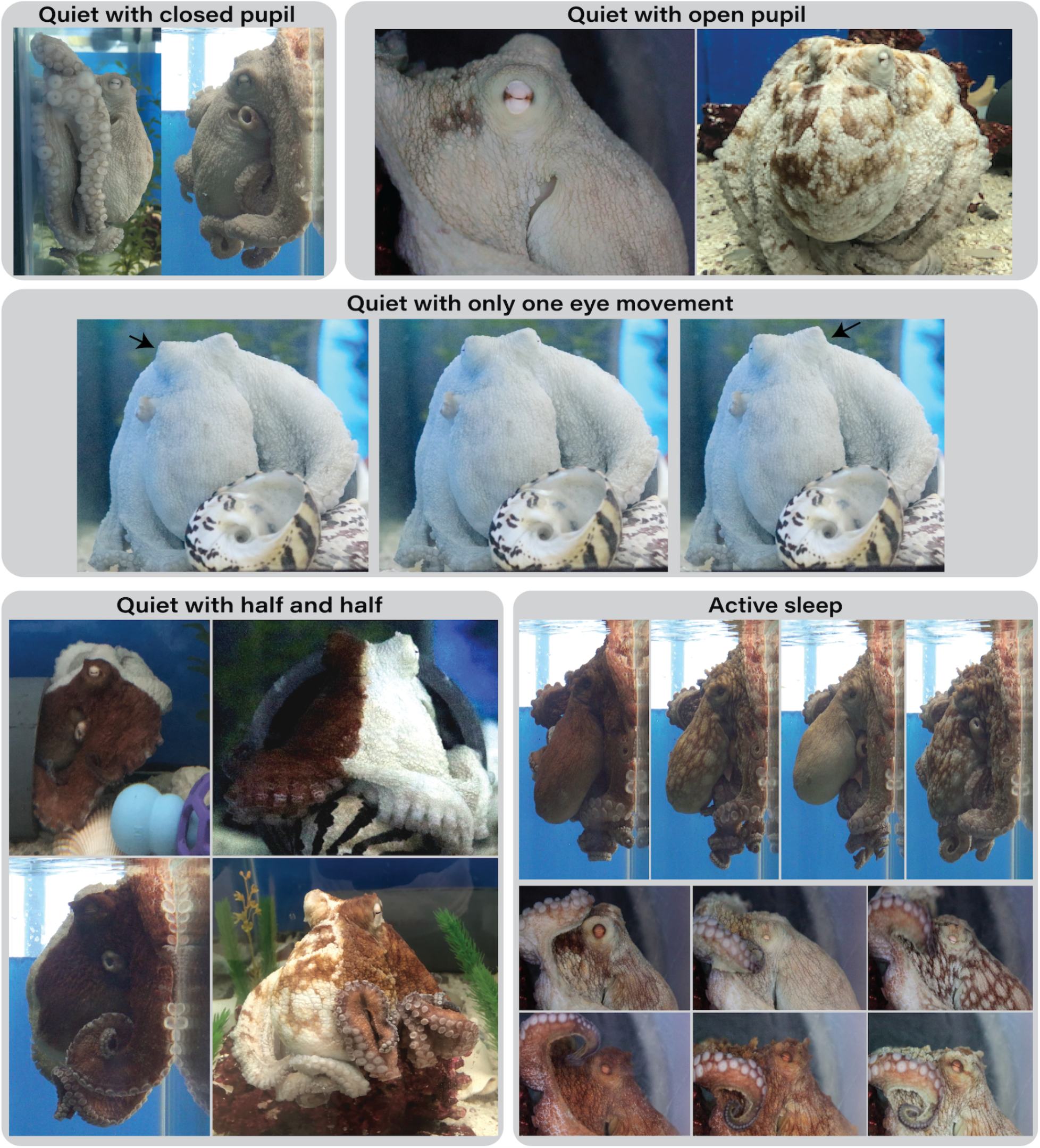
Images of *Octopus insularis* in the different quiescent states defined in Table 1, described by Medeiros et al. (2021).

All videos were reviewed in QuickTime Player 10 at least twice by two trained observers (EAR, MS) using all event sampling of our predefined behaviors. Any instances in which the octopus displayed one or more of the three behaviors were reviewed to determine if they qualified as abnormal episodes. We used standard definitions of octopus body parts, anatomy, and body patterns to provide detailed descriptions of its movements during these episodes (Mather & Alupay, 2016). Arms were labeled as L1 to L4 (L1 was the first arm to the left of the anterior midline of the animal going counterclockwise) and R1 to R4 (R1 was the first arm to the right of the anterior midline to the right going clockwise) (Leite & Mather, 2008).

## RESULTS

### Characteristics of abnormal episodes

To identify the occurrence of abnormal behavioral episodes, we reviewed a total of 3,600 hrs of video footage from three cameras gathered over 52 days from 11 Feb 2021 till the time of the octopus’s death on 04 Apr 2021. Four episodes fit our definition of abnormal episodes distinct from the animal’s normal behavior. In all four episodes, the octopus displayed a sequence of behaviors starting with abrupt emergence from quiescence or sleep, beginning with color changes and erratic and dysregulated movement of the arms and body, followed by the display of various evasive and aggressive behaviors generally associated with predator evasion and conspecific aggression in the wild (Figure 3; Supplementary Video S2).

Episodes 1,2 and 4 were brief in duration (mean = 44 s; *SD* = 7.6; range: 8–23 s) while Episode 3 was significantly longer (769 s). Abrupt movements, color changes, and dysregulated arm movements were also brief in Episodes 1, 2, and 4 compared to Episode 3 (Figure 2A-C & Figure 3). These movements of the body and apparent involuntary movement of the arms were not observed in other contexts (Figure 2 & Figure 3). Typically, the animal abruptly emerged from a sleep-like state, with its entire body rapidly changed to a solid dark coloration, sometimes alternating to mottled and back to solid red (Figure 1). Each of its arms appear to constrict and move erratically in different directions, some tightening while curled and other extended lengthwise upward and away from the animal (Figure 2 & Figure 3). In all episodes, dysregulated and erratic movements of the arms and body gave way to a sequence of fast movements and the display of various antipredator behaviors.

**Figure 2.**
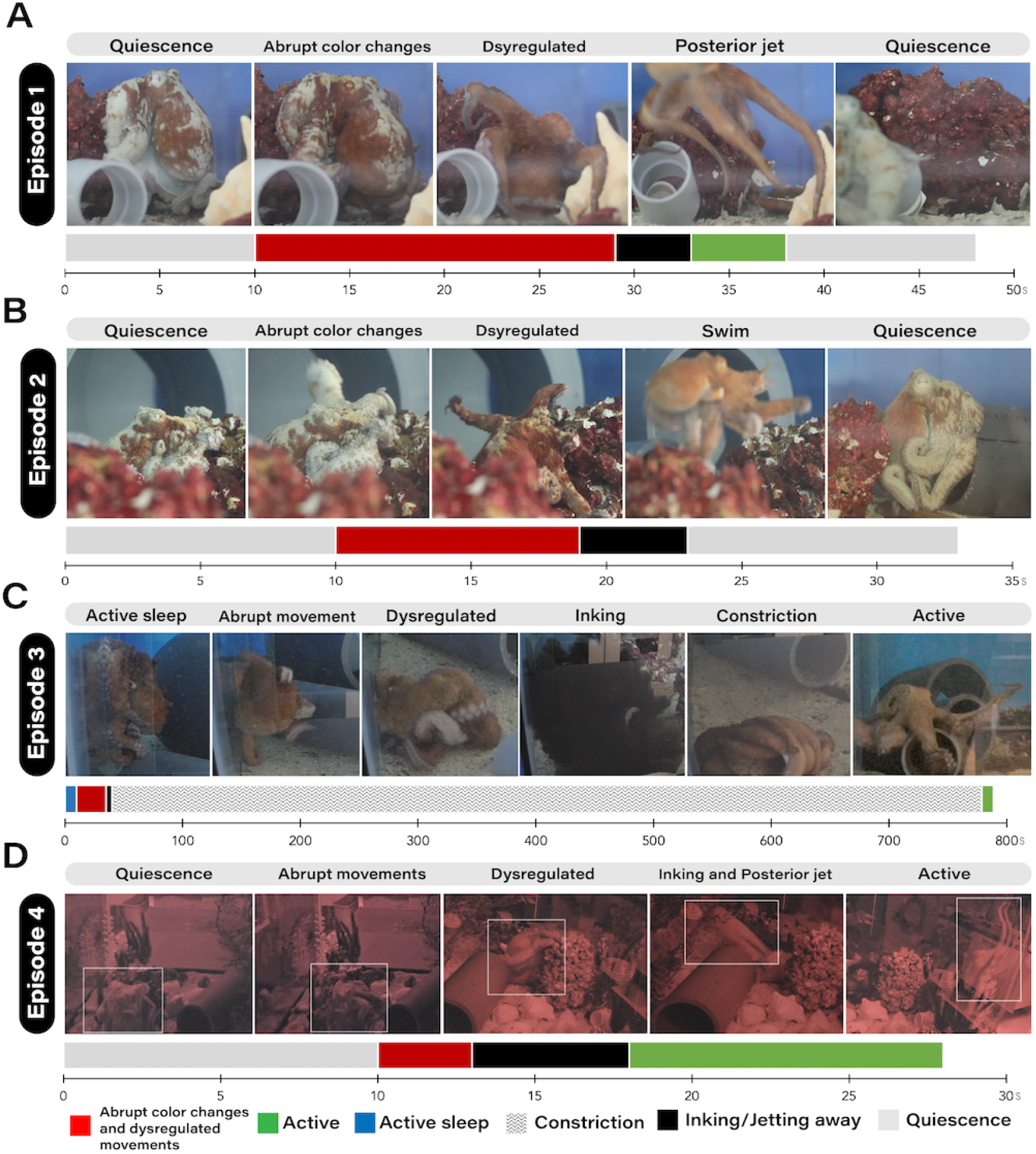
Graphic of behavioral sequences during abnormal behavioral events in a male *Octopus insularis*. Activity budgets of the octopus during each abnormal episode are displayed below each image sequence.

**Figure 3.**
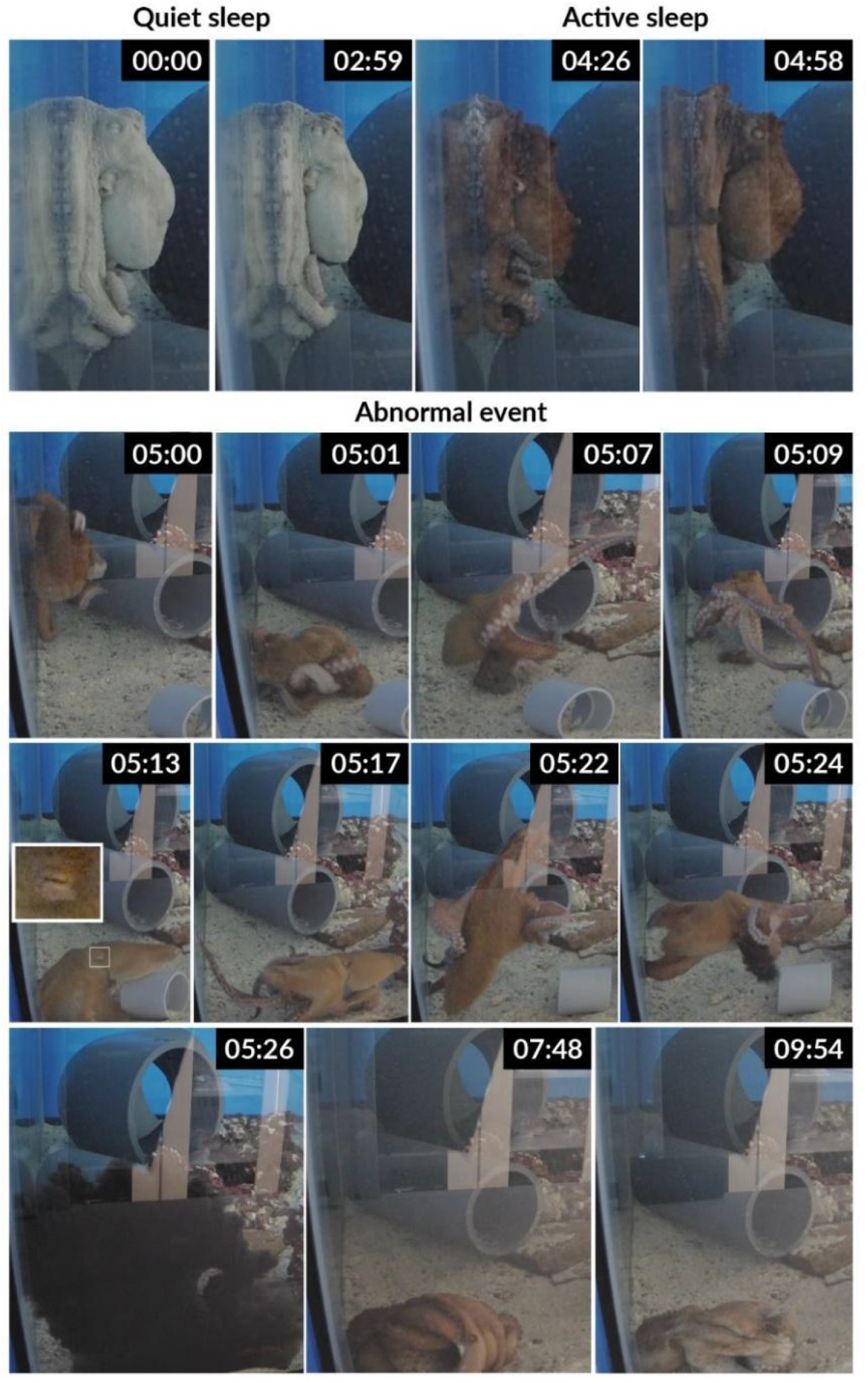
Sequence of images of abnormal sleep-associated behaviors in a male *Octopus insularis* in Episode 3. The octopus was largely in quiet sleep with eyes closed before the episode. Following a transition to active sleep, he abruptly swam away from his resting position with multiple arms writhing and appearing to spasm and engaged in a sequence of antipredatory and predatory positions as described in the text.

In the shorter-duration Episodes 1 and 2, the animal quickly returned to its previous behavior state of inactivity and quiescence. In Episodes 3 and 4, the animal displayed a variety of antipredator evasion behaviors and predatory behaviors associated with prey capture (Figure 2A-D & Figure 3). Please refer to the Supplementary Movie 1 for videos of the episodes. The episodes are numbered chronologically, but the major and clearest episode was #3.

### Descriptions of episodes and behavioral context of occurrence

#### Episode 1 - 2021-02-20 14:04:07

In Episode 1, the octopus had been in a quiescent state with mottled patterning on a red rock (Figure 2A) for the previous 15 minutes. He abruptly began to change color similarly to active sleep, with its eye open and turning yellow and exophthalmos of the visible left eye, and dark red across the rest of its surface. It then abruptly began moving its arms in a frantic fashion for curling its arms back slightly over its mantle while moving off the rock swimming backwards and upwards. He dragged a piece of PVC with his arms and rested on it, returning to quiescence.

#### Episode 2 - 2021-02-22 11:41:07 EST

The octopus had been stationary in a quiescent state with its pupils open on one of the dark red reef rocks (Figure 2B) for over 10 minutes. It then abruptly changed its color to dark red, eyes to yellow, and began moving its arms and body in a dysregulated fashion, appearing to lack control of its movements while curling its arms backward over its mantle and moving back and down from the rock towards the sandy substrate.

#### Episode 3 - 2021-02-23 11:25:01 EST

In Episode 3, the octopus had been in a quiescent state with the typical white appearance on the back wall of the tank, pupils narrowed to a slit (closed), and slight movements of the dangling arms (Figure 2C & Figure 3) for about 30 minutes. Small color changes were evident on the anterior of the mantle near the closed eyes. At 11:24:05, it entered active sleep: the entire body of the animal began to change dynamically in color and texture with movements of its visible eye, sucker contractions, and seemingly erratic or agitated movements of the arms and body (Figure 3). This behavior continued with numerous full-body color changes for 56 seconds as he assumed a dark red coloration. He then abruptly swam diagonally down along the wall, with his arms curled backwards over his body in a stereotypical posture observed while defending from or attacking eels (Hanlon & Messenger, 2017 page 131, also see Supplementary Figure S1) while moving his entire body at a 45 degree angle towards the left bottom of the tank (Figure 3). The animal spun around several times in a clockwise direction while near the bottom. The arms of its right side were curled, and the left arms were extended. Its mantle erected lengthwise, appearing pointed and elevated. The animal continued to spin with its arms moving frantically and its body positioned in various directions while still moving its body and multiple arms erratically Its eyes were both open at the time of the spasms. It then began to eject ink, filling half of the tank. As the ink began to clear, the animal was on its right side on the bottom tightly constricting a white PVC coupling (7 cm diameter, 6 cm length). Its arms were wrapped around the sides of the coupling and through the hollow portion. It remained solid dark red for 8 min before changing to mottled and camouflaged color. Now upright and alert, he continued to grasp onto the coupling as in preparation and consumption of prey with its mantle and web enveloping it in a ‘webover’ posture (Mather & Alupay, 2016). After 4 min largely stationary in consumptive behavioral postures, he began to move posteriorly across the tank, carrying the PVC coupling with him 10 cm across the tank. He resumed apparent normal activity following the episode, feeding successfully, and displaying normal baseline behaviors in the following days and weeks

#### Episode 4 - 2021-03-30, 23:50:17 EST

Episode 4 occurred at nighttime and our records are through the infrared camera. The octopus had been in a quiescent state or inactive on a rock (Figure 2D) for several minutes. It then displayed abrupt color changes and dysregulated movements for 3 seconds before inking and jetting away for 10 seconds, then inking a second time. It then began to move actively around the tank in Active state. The episode was registered in deep red illumination during the night.

## DISCUSSION

Understanding abnormal sleep-associated behaviors in invertebrates can provide insights into the nature of their sleep. We documented four episodes in an adult male *O. insularis* during which the animals displayed dysregulated movements of the arms and body, including violent thrashing in many directions in the longest episode. The dysregulated movements of the arms and body were accompanied by color changes to solid dark red immediately followed by evasive, antipredator defense behaviors. The occurrence of these episodes exclusively during sleep and quiescent states suggest they are associated with the sleep-wake cycle in this species. Behavioral displays associated with predator evasion and defense following body and arms spasms suggest that the octopus was in momentary to prolonged distress during abnormal sleep-associated episodes. We speculate that the complex behavioral sequences displayed in these episodes suggest octopuses experience parasomnias which may include nightmares with the potential to disrupt their sleep. If this were to be the case, this remarkable convergence of sleep behaviors in vertebrates and cephalopods points to the evolution of complex sleep in cognitively complex organisms with elaborate neural architectures.

Several hypotheses can be proposed to explain the observed episodes in the octopus. The behavior documented in our study differs significantly from any previously recorded instances. A notable pattern emerged, consisting of sudden awakenings followed by disorganized, erratic movements, paralleling waking antipredator responses typically seen in octopuses. This pattern, especially evident in Episode 3, could potentially represent dream-enactment behaviors. In the realm of human sleep, dream-enactment behaviors often involve individuals acting out occasionally violent dream scenarios during both REM and non-REM sleep, often in association with parasomnias like REM Sleep Behaviour Disorder (Nielsen et al., 2009; Ohayon & Schenck, 2010). While the existence of dreams in animals remains speculative, the complex movement behaviors observed during feline sleep lend credence to the possibility of dream-enactment. Cats with bilateral lesions of the locus coeruleus or the pontine pathway exhibited no differences in waking behavior or Slow Wave Sleep (SWS). However, during REM sleep, the typical muscle atonia associated with this stage was absent. Consequently, cats demonstrated oneiric behaviors: complex, stereotyped motor responses and sequences of behaviors, including actions reminiscent of prey stalking, defensive and aggressive behaviors (e.g., arched back, piloerection), flight behaviors, and grooming (Jouvet, 1979; Jouvet, 1962; Jouvet & Michel, 1959). These behaviors have been postulated to be outward manifestations of dream activity.

For the behaviors we observed to be interpreted as possible dream-enactment, there would need to be some ecological relevance to the sequence of behaviors we observed to the waking life of this octopus (Malinowski et al., 2021). In keeping with this, the behaviors we observed are consistent with the waking activity of octopuses in choice secondary defense behaviors outlined by Hanlon and Messenger (2008), where they describe the hierarchical structure cephalopods employ in deciding how to respond to threats. The primary defense behaviors include displays like deimatic and disruptive patterning with the intent of hiding from predators followed by protean behavior (unpredictable behaviors used to deter predators). Once a predator is detected, cephalopods typically engage in secondary defense, including inking, and jetting away from the area (Hanlon & Messenger, 2017). At the beginning of Episode 3, the octopus displayed behaviors consistent with protean responses to predators involving rapid and unpredictable color changes and shape changes (i.e., rapid neural polyphenism) during active sleep as described in the results. His initial abrupt movements involved backwards curling of the arms, consistent with behaviors observed in wild octopuses, in which they pull their arms tightly over their head and mantle leaving only the suckers exposed during antipredator displays to eels and when fighting conspecifics (MacGinitie & MacGinitie, 1968, Hanlon & Messenger 2017 p131, and Supplementary Figure). He then extended his mantle in an elongated and pointed shape with its body elevated and turned a solid red consistent with body patterns used in conspecific signaling and anti-predator defense in octopuses (Scheel et al., 2016; Mather & Alupay, 2016). This persistent dark coloration was observed most in the first few days of its arrival and afterwards was rarely seen in the animal’s daytime behavior, typically if it was startled by something outside of the tank (e.g., large moving objects). In Episode 3, the octopus displayed constricting behavior with its arms wrapped around a PVC coupling, an agonistic behavior observed between conspecifics involving at least one arm around the base of the mantle of another octopus (Huffard et al., 2010). This constriction behavior was reported in fatal constriction of a male *O. cyanea* by a female and in a non-fatal constriction of a *Wunderpus photogenicus* on a *Thaumoctopus mimicus* (Huffard & Bartick, 2015). This behavior has not been reported in *O. insularis*.

Besides dream-enactment, there are more complex behavioral and sleep disorders in which humans also act out complex, typically negative behaviors including sleep disturbances driven by sleep disorders like posttraumatic stress disorder (PTSD) (Germain, 2013; Campbell & Germain, 2016), or psychogenic nonepileptic seizures (PNES) observed in abnormal nocturnal sleep behaviors of humans (Pavlova et al., 2015), associated with distress to social or emotional stimuli indicating linkages with a variety of physiological and psychological processes (e.g., sensory, motor and psychological outputs). Nocturnal paroxysmal behavior and other parasomnias in humans often occur during transitions to and from sleep and result in the expression of behaviors indicating emotional and psychological distress. However, each of these alternatives require speculating a more complex disorder to justify the occurrence of these behaviors. Additionally, despite the similarities in cyclical alternation of ultradian sleep cycles between cephalopods and humans (e.g., Iglesias et al. 2017; Medeiros et al. 2021), the phylogenetic distance between cephalopods and humans and the vast differences in neurophysiology make it unlikely these episodes are analogous in physiological origin.

The behaviors displayed in each episode suggest the animal was in temporary distress, and absent any obvious external stimulus, that it may have been responding to internal stimuli or memories of previously experienced pain or trauma or long-lasting physiological damage. This specimen was wild caught, and nothing is known about its previous experiences. The animal arrived at the laboratory with evidence of having lost at least 60% or two different anterior arms (R1, L1) and damage to the distal 15% of a posterior arm (R4), most likely to predator attack. Pain experiences of the peripheral nerves in the arms (such as those incurred during the loss of arms to predators) are transmitted through central projections to the central nervous system within the mantle (Crook et al., 2013) and can result in long-term behavioral and neural hypersensitivity (Alupay et al., 2014). Octopuses subjected to injury in distal regions of the arms display spontaneous evasive behaviors and inking (Alupay et al., 2014) and localized pain responses, evidence of affective pain experience (Crook et al., 2021). The effects of these experiences can long-lasting impacts on octopus behavior, for example, Bock’s pygmy octopuses (*O. bocki*) injected with a painful noxious stimulus (acetic acid) displayed strong conditioned place avoidance to an initially preferred location and displayed spontaneous pain-associated behaviors, i.e., removal of its skin at the site of noxious chemical injection (Crook et al., 2021).

Predator attacks are likely traumatic to an octopus and several lines of evidence suggest long-term sensitization to predator encounters and sublethal injury function to improve strategies for avoiding future predation (Walters, 1994). For example, minor injury in the squid *Doryteuthis pealei* results in long-term sensitization in their behavioral and neuronal responses, leading injured squid to display sequences of antipredator behaviors faster than uninjured squid (Crook et al., 2013). Brief treatment of the injured squid with an anesthetic blocking nociceptive sensitization (i.e., pain responses) removed their early defensive responses and reduced their ability to escape and survival, supporting the notion the sensitization caused by predatory attacks drives improved responses to threats and increases chances of survival (Crook et al., 2013). Additionally, recuperating from limb tissue damage incurred during predation events (Lange, 1920; Voight, 1992) is energetically costly, and can require periods of inactivity, limiting their foraging efficiency, and/or may increase their vulnerability to predation (Wells & Clarke, 1996). Coupled with their capacity for long-term potentiation of memories (Hochner et al., 2003), escaped predation events and traumatic injuries likely contribute to long-term memories of specific negative experiences. For example, squid (*Euprymna scolopes*) showed lifelong neural excitability, cognitive deficits, and altered defensive behaviors in response to injury to the peripheral nervous system in early life (Howard et al., 2019).

The drivers for these episodes are unknown and we can only speculate as to what caused the behavior observed in the octopus. The first three episodes occurred within the first few weeks of its acclimation to the new environment, a period in which the animal often displayed antipredator body displays and increased arousal in response to the novel laboratory environment. These episodes could be the result of a naturally heightened arousal of octopuses during sleep in response to novel sensory stimuli.

Alternatively, it is possible that an imbalance in its light/dark cycle relative to its previous environment in the wild, or imbalances in oxygenation of its tank increased the likelihood of sleep disturbances. However, at this stage this assertion is largely speculative. Disturbance of the normal 12 hr/12 hr light/dark cycle of octopuses has been shown to cause a rebound in sleep (Meisel et al., 2011). However, at the time of all episodes there were no detectable stimuli present in its environment or the laboratory that would reasonably warrant the intensity of reactions observed. Differences in the life history of individual cephalopods are likely to influence their personality and responsivity to the external environments (Mather & Anderson, 1993).

Alternatively, during senescence, octopuses undergo degeneration of body and nervous system function and cognitive declines that could contribute to disturbed sleep (Anderson et al., 2002; Halm et al., 2000; Holst & Miller-Morgan, 2020). Early in senescence, captive giant Pacific octopuses (*Enteroctopus dofleini*) display a behavioral hypersensitivity in which mild dermal stimulation could be perceived as aversive (Holst et al., 2022). Given our octopus had the most episodes and most severe episode (Episode 3) nearly 6 weeks prior to its death, these behaviors could indicate it was experiencing effects of senescence yet to be detected in cephalopods.

Our report is limited in scope given it focuses on one individual. How common these episodes may be in octopuses is unknown. A recent study that analyzed hundreds of hours of octopus behavior during active and inactive states did not detect abnormal sleep-associated episodes in *O. insularis* (Medeiros et al., 2021). These rare and brief events were only captured due to our extensive monitoring with high-resolution video cameras, enabling the documentation of the octopus’s activity throughout its life. The infrequency of these episodes and their brief duration make it unlikely for these behaviors to be detected in wild animals where they are often out of sight. *O. vulgaris* and *O. insularis* in the wild protect themselves from predation by living in their dens. An octopus transitioning from stationary inactivity to abrupt color changes and movement like the behaviors we observed would dramatically increase the probability of their detection by nearby predators and increase their risk of mortality.

Future studies should carefully document the 24 hr waking and sleeping behaviors of multiple octopuses of different species to explore how their activity cycles influence sleep and the possible occurrence of dream-enactment and/or parasomnias. Our findings support the use of 24 hr continuous video monitoring with cephalopods in laboratory settings as a powerful means of identifying these rare behavioral episodes. Cameras positioned on both sides of the tank would help to reduce out-of-sight periods when an octopus can be obscured by objects in its enclosure (e.g., PVC tubes, rocks). Tests of dream-enactment in captive octopuses could involve cognitive experiments focused on memory and learning and the development of underwater electroencephalogram sensors to identify if active sleep and other sleep stages reflect waking neural activity and body patterns (Malinowski et al., 2021). If detected in captivity as in our case, detailed necropsies of afflicted individuals are needed to identify possible linkages of these behaviors to neurological damage or disorders. Capturing the sleep behavior of wild octopuses could employ artificial homes placed in their habitats rigged with sufficient lighting, or temporary captures or octopuses.

## CONCLUSIONS

In this study we present novel evidence for abnormal sleep-associated behavioral episodes in *O. insularis*. The antipredator and agonistic behaviors observed during two episodes suggests the animal was in distress during these occurrences. The behavioral sequences displayed by this octopus upon emerging from disturbed sleep were similar to behavioral responses to nightmares, night terrors, and other parasomnias in humans, with a narrative structure resembling waking defense behaviors in octopuses. We should caution the reader that because of the single-subject nature of this study, we cannot rule out specific neurological abnormalities of our octopus or other issues; while we have done our best to show that the baselines were normal, this study cannot be considered conclusive until replicated. The potential occurrence of this phenomenon in octopuses would shed an unexpected light on the convergent evolution of sleep in distantly related organisms with complex neural architectures.

## ACKNOWLEDGMENTS

Thanks to Brett Grasse at 8 Arm Assistance LLC Advisory Services for his expert consultation in setting up the octopus enclosures. Thank you to Felix Miranda at Best Aquarium Service NYC for his invaluable support in establishing and upkeeping the octopus laboratory. Thanks to David Scheel at Alaska Pacific University for his insights on octopus behavior and sleep. Thanks to Connor Gibbons at the Axel Lab (Zuckerman Institute, Columbia University) for his assistance with octopus keeping and enclosure improvements. Thanks to the veterinary team at the comparative Bioscience Center at The Rockefeller University. This research was conducted at the Laboratory of Integrative Neuroscience and Comparative Bioscience Center at The Rockefeller University. All work was approved by the Institutional Animal Care and Use Committee (IACUC) at The Rockefeller University. While the manuscript was being prepared, Leenoy Meshulam kindly made us aware of their own paper in advance of publication (Pophale, Shimizu et al, 2023) where changes in electrical activity can be observed at the onset of active sleep.

## Supplementary Information

**Supplementary Video (attached separately)**. Video compilation of the four abnormal sleep-associated episodes documented in a male *Octopus insularis*.

**Supplementary Figure.**
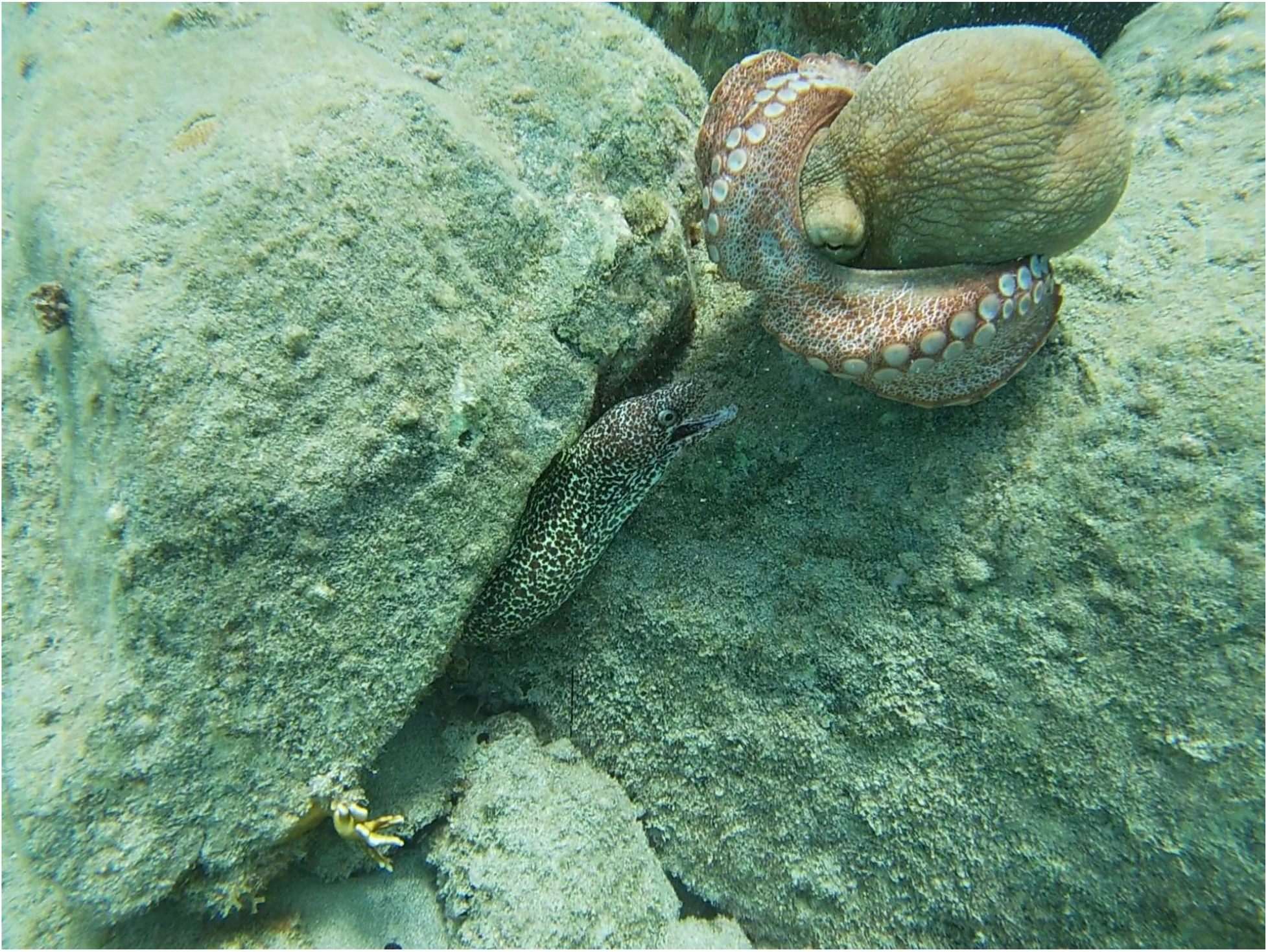
An octopus (apparently *O. insularis*) in confrontation with a spotted eel, adopting a stereotypical posture crossing its arms behind its back to prevent the tips from being bitten (Hanlon and Messenger, 2017 page 57?). Recorded in St. John, USVI. Courtesy of Mitch Helwig (ig:@mitchyonstj), all rights reserved.

